# Discovery of UFO Proteins: Human-Virus Chimeric Proteins Generated During Influenza Virus Infection

**DOI:** 10.1101/597617

**Authors:** Yixuan Ma, Matthew Angel, Guojun Wang, Jessica Sook Yuin Ho, Nan Zhao, Justine Noel, Natasha Moshkina, James Gibbs, Jiajie Wei, Brad Rosenberg, Jeffrey Johnson, Max Chang, Zuleyma Peralta, Nevan Krogan, Christopher Benner, Harm van Bakel, Marta Łuksza, Benjamin D. Greenbaum, Emily R. Miraldi, Adolfo Garcìa-Sastre, Jonathan W. Yewdell, Ivan Marazzi

**Affiliations:** Department of Microbiology, Icahn School of Medicine at Mount Sinai, New York, NY 10029, USA; Laboratory of Viral Diseases, National Institute of Allergy and Infectious Diseases, NIH, Bethesda, MD 20892, USA; Department of Cellular and Molecular Pharmacology, University of California, San Francisco, San Francisco, CA 94158, USA; Department of Medicine, School of Medicine, University of California San Diego, La Jolla, CA 92037, USA; Department of Genetics and Genomic Sciences, Icahn School of Medicine at Mount Sinai, New York, NY 10029, USA; Tisch Cancer Institute, Icahn School of Medicine at Mount Sinai, New York, NY 10029, USA; Department of Medicine, Hematology and Medical Oncology, Icahn School of Medicine at Mount Sinai, New York, NY 10029, USA; Department of Oncological Sciences, Icahn School of Medicine at Mount Sinai, New York, NY 10029, USA; Department of Pathology, Icahn School of Medicine at Mount Sinai, New York, NY 10029, USA; Divisions of Immunobiology and Biomedical Informatics, Cincinnati Children’s Hospital, Cincinnati, OH 45229, USA; Department of Pediatrics, University of Cincinnati College of Medicine, Cincinnati, OH 45257, USA; Department of Microbiology, Icahn School of Medicine at Mount Sinai, New York, NY 10029, USA; Global Health and Emerging Pathogens Institute, Icahn School of Medicine at Mount Sinai, New York, NY 10029, USA; Division of Infectious Diseases, Department of Medicine, Icahn School of Medicine at Mount Sinai, New York, NY 10029, USA; Department of Microbiology, Icahn School of Medicine at Mount Sinai, New York, NY 10029, USA; Global Health and Emerging Pathogens Institute, Icahn School of Medicine at Mount Sinai, New York, NY 10029, USA; The State Key Laboratory of Reproductive Regulation and Breeding of Grassland Livestock, College of Life Sciences, Inner Mongolia University, Hohhot, 010070, China

**Keywords:** Influenza, uORFs, viral evolution, chimeric protein

## Abstract

Influenza A virus (IAV) is a threat to mankind because it generates yearly epidemics and poorly predictable sporadic pandemics with catastrophic potential. IAV has a small RNA genome composed of 8 mini-chromosomes (segments) that constitute a 5’UTR followed by a coding region and a 3’UTR. Transcription of IAV RNA into mRNA depends on host mRNA, as the viral polymerase cleaves 5’m7G-capped nascent transcripts to use as primers to initiate viral mRNA synthesis. We hypothesized that captured host transcripts bearing AUG could drive the expression of upstream ORFs in the viral segments, a phenomenon that would depend on the translatability of the viral 5’UTRs. Here we report the existence of this mechanism, which generates host-virus chimeric proteins. We label these proteins as <u>U</u>pstream <u>F</u>lu <u>O</u>RFs (UFO). Depending on the frame, two types of host-virus UFO proteins are made: canonical viral proteins with human-derived N term extensions or novel uncharacterized proteins. Here we show that both types are made during IAV infection. Sequences that enable chimeric protein synthesis are conserved across IAV strains, indicating that selection allowed the expansion of the proteome diversity of IAV in infected cells to include multiple human-virus proteins.

## INTRODUCTION

Influenza A virus (IAV), of the family *Orthomyxoviridae*, is a highly contagious human and animal pathogen responsible for significant levels of morbidity and mortality worldwide. The virus bears a single-stranded, negative-sense RNA genome that is organized into eight segments (Bouvier and Palese, 2008). Viral mRNA transcription and genome replication both occur within the host nucleus, and require the three-subunit viral RNA-dependent RNA polymerase (RdRP) complex comprising of PB1, PB2 and PA proteins (Bouvier and Palese, 2008; Te Velthuis and Fodor, 2016).

IAV viral mRNA synthesis is primed using 5’ methyl-7-guanosine (m7G) capped short RNA sequences cleaved from host RNA polymerase II (RNAPII) dependent transcripts. During this process, named “cap-snatching”, PA cleaves host-capped RNA bound to PB2 to generate 7-20 nucleotide long, capped RNA fragments (Dias et al., 2009). These host-derived fragments are then utilized by PB1 to initiate the transcription of viral mRNAs (Plotch et al., 1981; Reich et al., 2014). Consequently, IAV viral mRNAs are not only genetic hybrids that include both host and viral derived sequences, but also possess diverse 5’ sequence heterogeneity (Koppstein et al., 2015; Sikora et al., 2017). Once made, viral mRNA is exported to the cytoplasm and translated by the host machinery.

Each segment of the IAV genome encodes one major open reading frame (ORF) flanked by 5’ and 3’ untranslated regions (UTRs). The IAV segments code for eight major structural and non-structural proteins (PB2, PB1, PA, HA, NA, NP, M1, NS1). In addition, IAV utilizes several different mechanisms to expand the coding capacity of the individual segments to generate additional proteins. Segments 7 and 8 are spliced to produce M2 and NEP proteins, respectively (Inglis and Brown, 1981; Lamb and Lai, 1980; Lamb et al., 1981). PB1-F2 and N40 (Chen et al., 2001; Wise et al., 2009) proteins arise from leaky ribosomal scanning of IAV segment 2. Segment 3 encodes an alternative protein, PA-X, generated by +1-ribosomal frameshift during the translation of PA protein (Jagger et al., 2012). Segment 3 also produces several N-terminally truncated forms of PA due to alternate start codon (AUG) usage (Muramoto et al., 2013). Additional viral proteins, such as M42, might be encoded from alternative spliced mRNAs (Wise et al., 2012).

Intriguingly, full genome studies on IAV isolates have revealed that the length and sequence context of these accessory proteins varies between IAV strains. These differences are often correlated with altered virulence and/or responses of host cells. For example, PB1-F2 protein derived from the IAV laboratory strain A/H1N1/Puerto Rico/8/1934 induces apoptosis in host cells through the interaction with BAK/BAX (Chen et al., 2001). In contrast, PB1-F2 from a H5N1 IAV strain (A/H5N1/Hong Kong/156/1997) is non-mitochondrial and not pro-apoptotic (Chen et al., 2010). As such, identification of novel accessory proteins and the breadth of their diversity across different strains of IAV may provide insight into viral replication and the interplay with the infected host.

In this manuscript, we describe the existence of IAV-human protein chimeras. We show that at least three such chimeric proteins are synthesized during influenza virus infection. These proteins are initiated from cap-snatched RNA sequences with upstream AUGs (uAUGs) that initiate translation of the IAV 5’ UTR and the downstream viral segment. Through this mechanism, host uAUGs create either viral protein N-terminal extensions and/or the synthesis of novel, heretofore uncharacterized host-viral proteins (UFOs). We show that both types of proteins are expressed in infected cells, as our analyses reveal the existence of HA and NP extensions driven by host RNA and also identify an uvORF in segment 2 that generates a novel, ∼77 amino acid long protein (PB1 <u>U</u>pstream <u>F</u>lu <u>O</u>RF; PB1-UFO). Full length PB1-UFO is conserved in more than 90% of isolates of IAV. HA and NP extensions are conserved in 99% of IAV isolates. Overall, our analysis reveal that host-viral protein chimeras are (1) segment-specific, (2) conserved across IAV strains and (3) undergo differential selection pressures according to 5’ UTR and coding region constrains, resulting in fixation of N term extensions and novel ORFs that are sampling evolutionary space through genomic overprinting.

## RESULTS

### IAV 5’ “UTRs” are potentially translatable

IAV transcription is initiated by host RNA cap snatching (**Figure 1A**). This process generates 5’ host derived extension of IAV segments. We hypothesized that this mechanism, used to express canonical viral proteins (**Figure 1B**, Outcome 1), could generate upstream host-virus chimeric ORFs with coding potential. Depending on the reading frame, an upstream host derived AUGs may either initiate the synthesis of N-terminal extended viral proteins (**Figure 1B**, Outcome 2) or novel uvORFs that overprint the canonical viral ORF (**Figure 1B**, Outcome 3). These outcomes are contingent on three assumptions: (1) premature stop codons are not present in translation frames of interest (2) viral “UTRs” lacking stop codons are evolutionarily conserved in IAV strains (3) AUGs are present in cap-snatched host sequences enabling translation of host-virus chimeric RNA.

**Figure 1.**
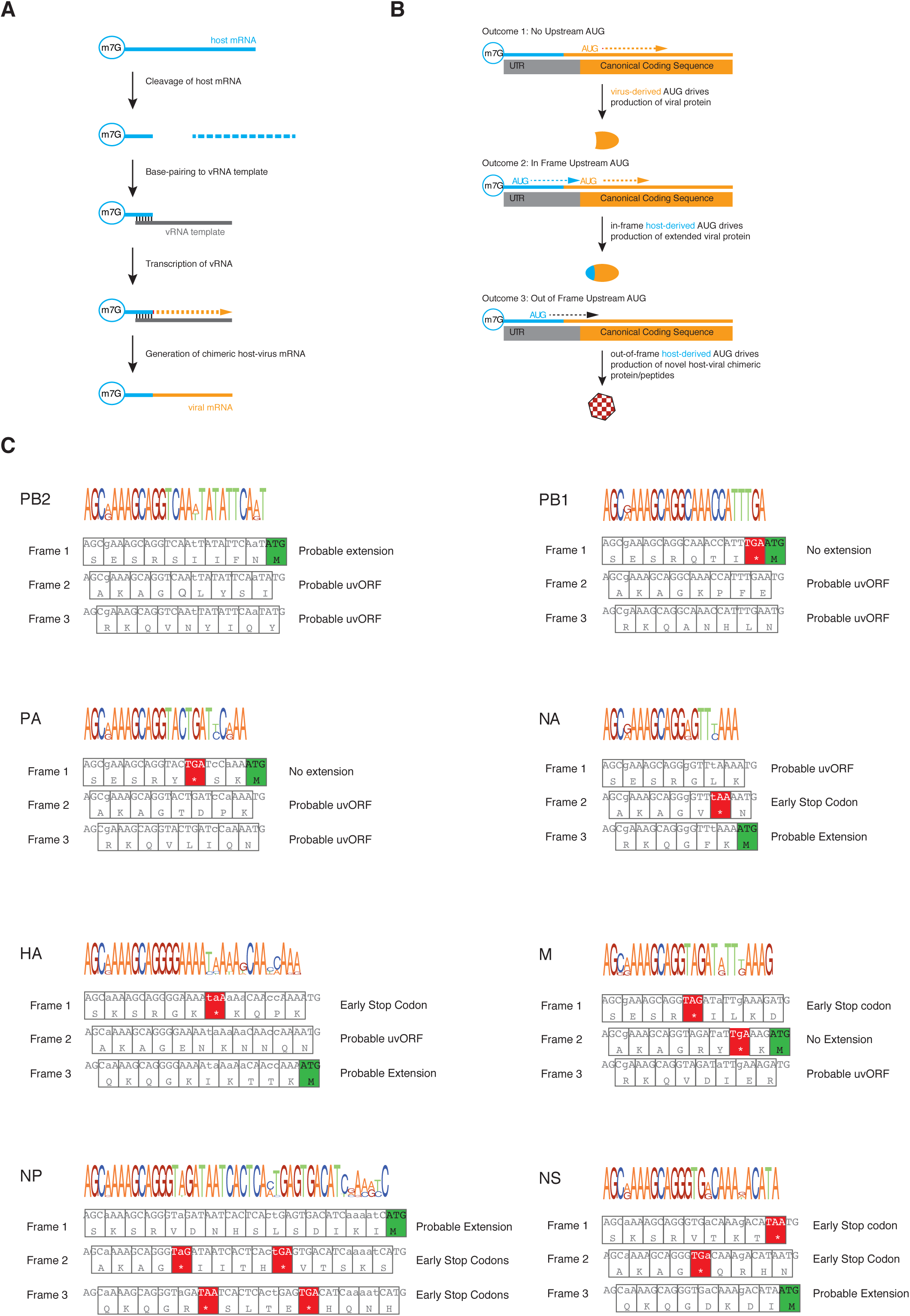
IAV 5’ UTR are conserved and translatable in all three reading frames. (A) Schematic of viral cap-snatching occurring during IAV infection. (B) Presence of upstream AUGs in host-derived segments of viral mRNA may drive the formation of viral protein extensions or novel host-viral chimeric proteins. (C; top panels) Predicted peptide sequences derived upon translation of all three ribosome reading frames in the 5’UTR. (C; lower panels) Sequence conservation analysis of IAV H1N1 strain 5’UTR within individual viral segments.

To address the first two points, we analyzed the nucleotide sequence variability within the 5’UTRs of all eight segments among all H1N1 strains available from the GISAID Database (Shu and McCauley, 2017). 5’UTRs are highly conserved within each individual segment, as shown by the positional weight matrices (**Figure 1C**, top panels and **Figure S1**). To determine if viral 5’UTRs can be translated to generate long peptides, we retrieved the most commonly occurring 5’UTR nucleotide sequences per segment (**Figure S1**). These sequences were then translated in all three strains *in silico* (**Figure 1C**). This revealed that the UTRs of 3 (PB1, PA and M) of 8 viral segments possess conserved stop codons in-frame and upstream of the major ORF start codon. Thus, 5/8 viral segments have the potential to code for N-terminally extended viral proteins if an upstream AUG is captured from host mRNA. Surprisingly, we also detected the absence of stop codons in the alternate translation reading frames of several viral segments (**Figure 1C**, Segments PB2, PB1, PA, NA and HA). We were thus intrigued with the possibility that these segments encode novel long peptides given an upstream, host-donated AUG in the right context.

### Host RNA bearing upstream AUGs are present in viral mRNA

We next determined the abundance of AUGs in host-snatched sequences generated from PR8 infected-A549 cells by RNA-sequencing. Host oligonucleotides with AUG codons constituted approximately ∼12% of all cap-snatched sequences, and were present at similar ratios in all eight segments of the virus (**Figure 2A**).

**Figure 2.**
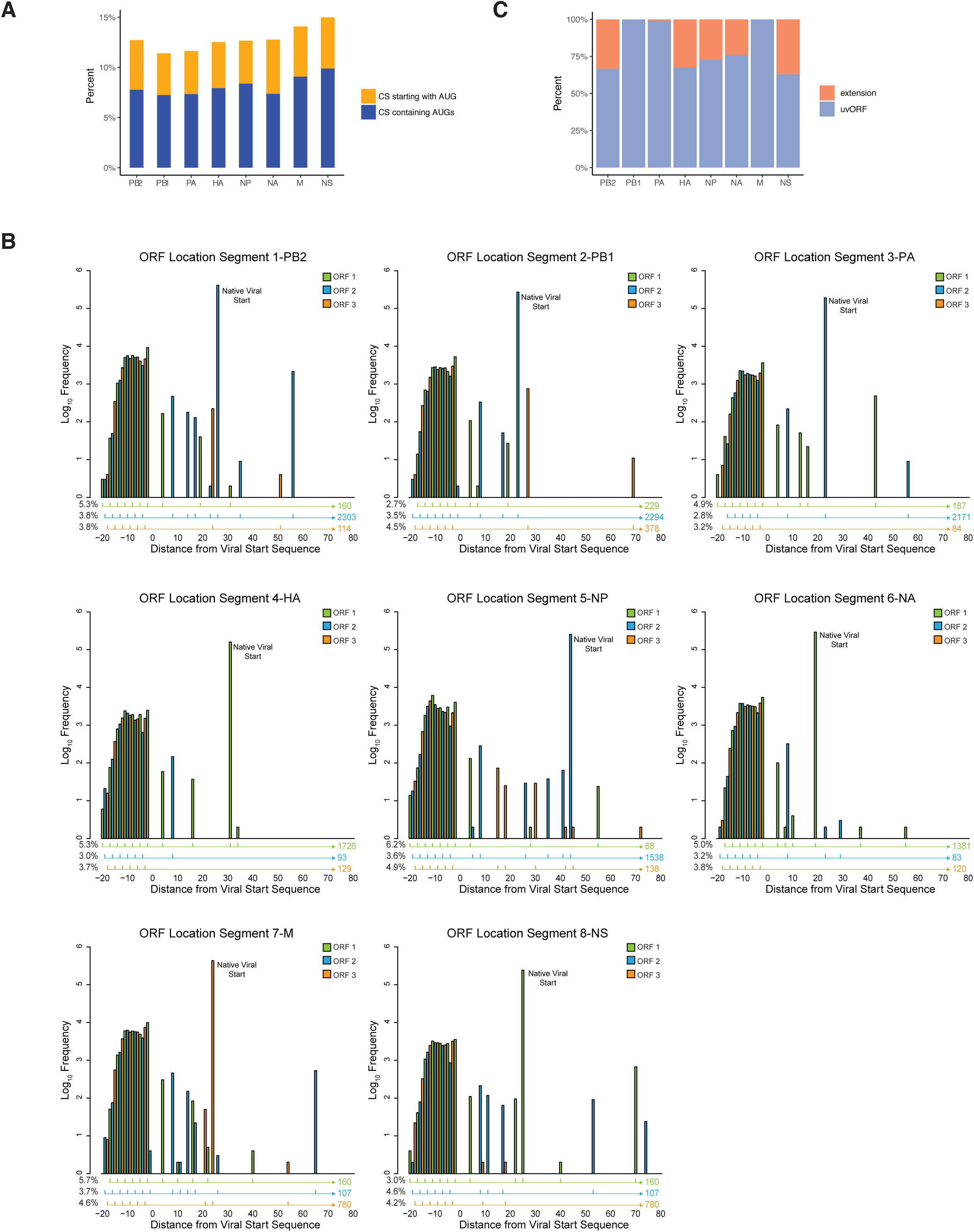
Upstream AUGs are present in host derived viral RNAs. (A) Percentage of cap snatched sequences bearing uAUGs in the first codon (yellow) and bearing uAUGs at other positions (blue). (B) Incorporation of host transcript sequences increases the diversity of putative alternative start codons. For each viral segment, the frequency and position of alternative start codons is shown relative to native start of the viral genes. For each reading frame, the frequency and location of the first in-frame stop codon are indicated. (C) Percentage of N-terminal protein extension (orange) and uvORFs (blue) derived from cap snatched sequences bearing uAUGs in the 8 viral segments.

Do uAUGs result in N-terminal extensions of viral proteins or generate uvORFs *in silico*? We aligned and extended viral derived sequences from sequenced host-virus RNA chimeras to match the reference sequences of the A/H1N1/Puerto Rico/8/1934 IAV. Viral UTRs with uAUGs were then translated *in silico* revealing all possible N-terminal protein extension and/or uvORFs in the data set. We define N-terminal protein extensions as ORFs with uAUGs in frame with the canonical ORFs and without a stop codon in the uORF. By contrast, we define ORFs with uAUGs out of frame with the canonical ORF as uvORFs. Putative sequences that would generate a novel ORF but would not contain a stop codon across the whole length of a viral segment were excluded because of the inherent instability of mRNA lacking stop codon (Simms et al., 2017). In an effort to be stringent with our analysis, host sequences that begun with an AUG at the 5’ were also removed from our analysis as it is unclear if the ribosome would be able to recognize these as start codons (**Figure 2A**, yellow bars). uvORF length filters were also not applied at this stage of the analysis as we reasoned that the ribosomal complex initiation and assembly should be independent of ORF length (Chew et al., 2016).

We mapped host-derived sequences in PR8 infected A549 via RNA sequencing. Our analysis revealed that host-derived uAUGs are present in all three translational reading frames, and at similar frequencies in the eight viral segments (**Figure 2B**). As expected (**Figure 1C and S1**), individual viral segments exhibited different propensities to generate N-terminally extended proteins (Orange bars; **Figure 2C**) compared to uvORFs (Blue bars; **Figure 2C**). ∼19% of uAUGs in host-derived sequences in PB2, HA, NP, NA and NS segments (**Figure 2C**) initiate N-terminal extensions of the major ORF, but next to none in PB1, PA and M segments. uvORFs are present in all segments at significant frequencies. Given that that viral genes are among the highest expressed RNA in the cells during infection, this suggests that uvORF containing viral RNAs are likely to be present at levels similar to most other host mRNA in the cell.

### Host-derived uAUGs drive the translation of uvORFs and N-terminal extensions during infection

We next sought to determine if viral N-terminal extensions or uvORFs are translated during infection. In silico analyses suggest that, as a function of the frame (F), the probable N-terminal extensions in PB2, HA, NP, NA, NS segments are very consistent in lengths within the given individual segment (**Figure 3A**). Extensions ranged from 9 – 17 amino acids, with the longest occurring in the NP segment (**Figure 3A**). In contrast, the lengths of uvORFs in the PB2, PB1, PA, HA and M segments hovered at or below 20 amino acids (**Figure 3B**). Most importantly, we found long conserved uvORFs in PB2, PB1, PA and HA segments, ranging from 40+ residues (HA) to nearly 80 residues (PB1).

**Figure 3.**
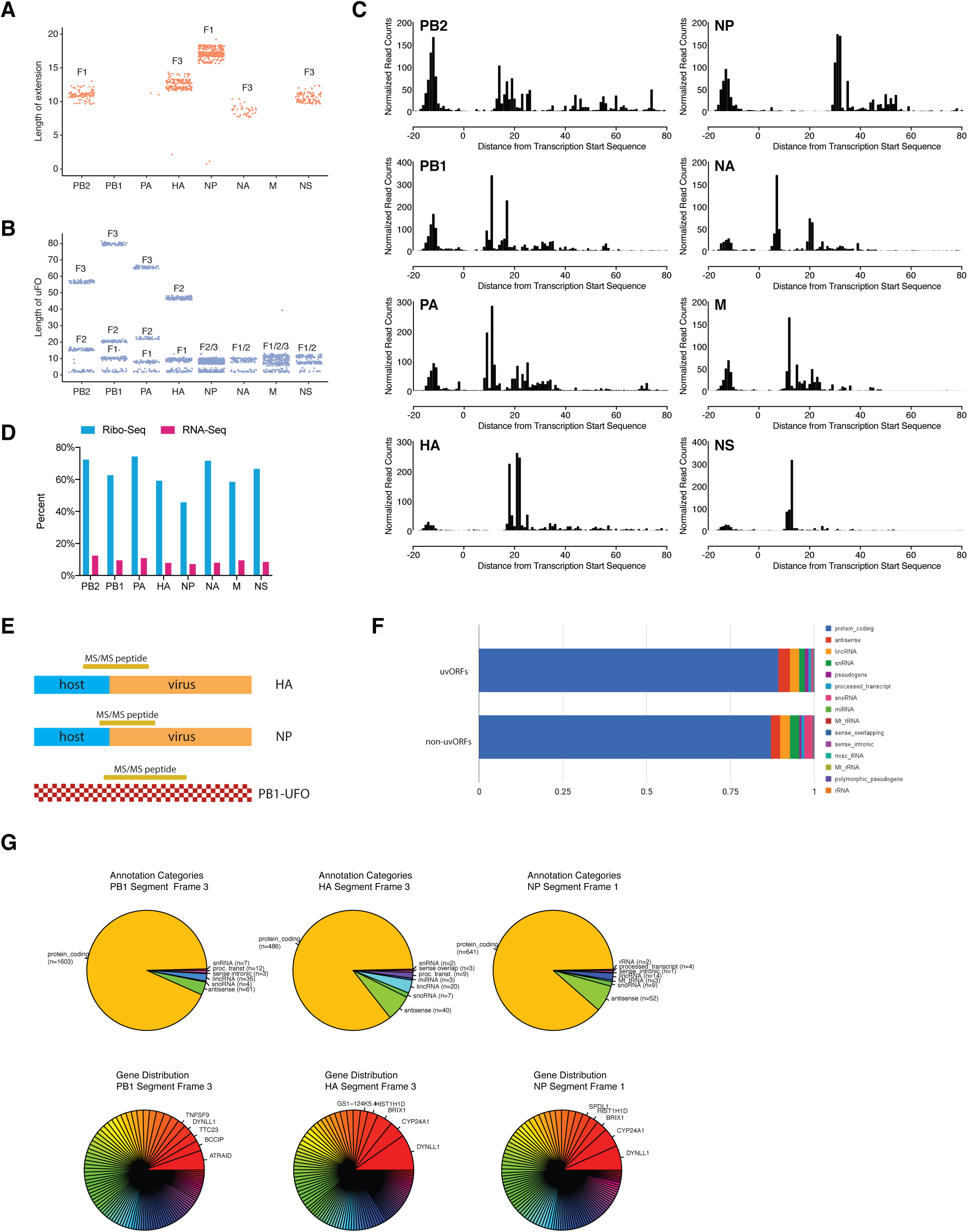
IAV 5’UTRs are translated. (A) Length distribution of N-terminal protein extension in individual segments. Each dot represents a protein predicted to be translated from a unique host–virus chimeric RNA. F represents the reading frame. (B) Length distribution of uvORFs of individual segments. Each dot represents a protein predicted to be translated from a unique host–virus chimeric RNA. F represents the reading frame. (C) Ribosome profiling of harringtonine-treated A549 cells infected with A/Puerto Rico/8/1934 (H1N1). (D) Frequency of primers containing AUG codons in harringtonine Ribo-seq and RNA-seq datasets. (E) Schematic shows peptides identified in HA, NP N-terminal extension and PB1-UFO via Mass Spectrometry analysis. (F) Source of CS events contributing to PB1-UFO. (G) Transcript biotypes contributing CS sequences for vmRNAs leading to uORFs and those that do not for segments PB1, HA and NP (upper panels). Relative CS abundance for the top-100 genes contributing to uORFs. Names and CS event read counts are shown for the top 5 genes (lower panels).

If snatched host uAUGs initiate translation of viral 5’UTRs, these sequences should be enriched in translating ribosomes in infected cells. Using RNA-seq and Ribo-seq, we mapped the 5’ end of ribosome footprint sequences from harringtonine-treated PR8-infected cells to the viral genome. This revealed an accumulation of reads ∼12nt upstream of the IAV canonical start codons in all eight segments (**Figure 3C**). Notably, we observed a large number of reads in the host-derived portion of the 5’ UTR, consistent with ribosome initiation. Furthermore, host sequences demonstrated a 7.5-fold enrichment in the Ribo-seq data set *vs.* the host primer sequences present in the RNA-seq data set of poly A containing IAV mRNA (**Figure 3D**).

If ribosomes initiate on host derived AUGs, many of the 5’ sequences will be too short to extend from the ribosome, given the brevity of their snatched caps, making P-site phasing problematic by standard Riboseq analysis. We therefore used the location of AUGs within the primers to identify the reading frame being translated. With few exceptions, initiation occurred evenly in all three reading frames. AUG codons tended to aggregate closer to the transcriptional start site, despite being depressed at the −4 position in all segments. Frequencies also tended to be lower towards the 5’ end of the primer (**Figure S2A**). This phenomenon is also observed when the frequency of primers containing AUG is compared to primer length **(Figure S2B)**.

Finally, to verify that chimeric proteins are indeed translated, using targeted proteomic analysis we evaluated the presence of UTR-derived and chimeric peptides in PR8 IAV infected cells. We unequivocally identified peptides that originate from predicted N-terminal extensions of the NP and HA and the long uvORF present in PB1 segment (**Figure 3E**).

Most host-derived sequences were from protein coding genes (**Figure 3F**), with similar distributions between the three segments (**Figure 3G, top panels**). Host caps were derived from different genes (**Figure 3G, bottom panels**) and were predominantly obtained from high expressing mRNAs (**Figure S2C**).

### PB1-UFO is a host-virus chimeric protein expressed during infection

We were especially intrigued by the ∼77-amino acid uvORF present in the 5’ UTR of IAV segment 2 (encoding PB1). This is one of the longest, conserved non-canonical ORFs in IAVs (**Figure 4A**). We designate this protein PB1-UFO and proceeded to characterize it.

**Figure 4.**
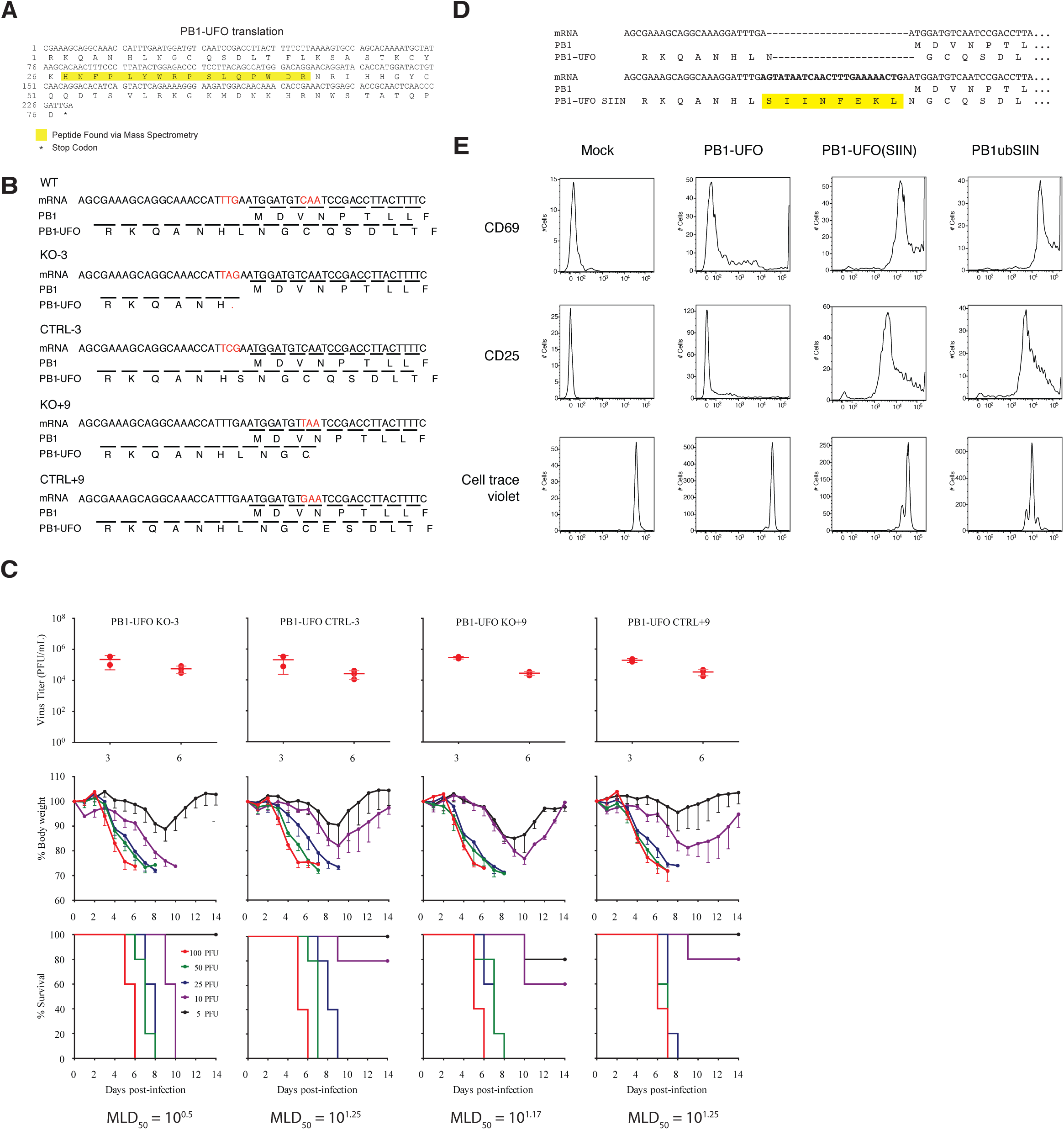
PB1-UFO. (A) Nucleotide and amino acid sequence of PB1-UFO protein. PB1-UFO peptide detected in mass spectrometry is highlighted in yellow. (B) Schematic of mutations used to construct PB1-UFO-null and control viruses in the PR8 virus background. (C) Viral titers (top panels), percentage of body weight loss (middle panels) and survival curves (lower panels) of mice infected with differing concentrations of the indicated viruses. MLD50 of each virus is indicated at the bottom. (D) Nucleotide and amino acid sequence of PB1 and PB1-UFO N-terminal sequences with the SIINFEKL peptide insertion in 5’ UTR to generate the PB1-UFO (SIIN) virus. (E) Infected DC2.4 co-cultured OT-I activation assay. CD69 and CD25 expression at 24 hours post co-culture, and cell proliferation assay at 48 hours post co-culture.

To evaluate the physiological role of PB1-UFO during infection we used reverse genetics to generate wild-type control (CTRL-3 and CTRL+9) and PB1-UFO-deficient mutant (KO-3 and KO+9) IAVs in the H1N1/Puerto Rico/8/1934 (PR8) strain background (**Figure 4B**). As PB1-UFO is predicted to translate from an alternative −2 reading frame, we could make single nucleotide substitutions to introduce premature stop codons for PB1-UFO without modifying the amino acid sequence of PB1 (**Figure 4B**). PB1-UFO truncating mutations (KO-3 and KO+9) were introduced at either the −3 or +9 nucleotides relative to the PB1 start ATG codon. We included viruses with mutations that did not disrupt either the PB1-UFO or PB1 reading as controls (CTRL-3 or CTRL+9). Mutant and control viruses all yielded stocks with similar particle counts as indicated by HA titers (**Figure S3A and S3B**). All viruses demonstrate similar growth in MDCK cells (**Figure S3A and S3B**) at high (40°C; **Figure S3A**) or physiological (37°C; **Figure S3B**) temperatures, demonstrating that PB1-UFO is not required for replication under these conditions.

PB1-UFO-deficient viruses were also able to replicate normally in the lungs of infected BALB/c mice as measured by virus yields after intranasal infection. Mice infected with PB1-UFO-deficient or control viruses displayed weight loss (**Figures 4C, middle panels**) over a range of infecting doses. Survival curves of mice infected with PB1-UFO-deficient viruses and controls revealed similar minimum lethal doses (MLD_50_) for both the KO-3 and KO+9 mutant viruses (**Figure 4C, column 1 and 3**) when compared to CTRL-3 and CTRL+9 viruses (**Figure 4C, column 2 and 4**). These data indicate that PB1-UFO is non-essential for virulence in mice, a phenomenon that is shared by most non-canonical protein from IAV. Since mutations in accessory IAV proteins, which cause subtle differences in pathogenesis, often display molecular phenotypes, we therefore isolated RNA from the lungs of mice infected with 100 PFUs of KO-3, CTRL-3, KO+9 or CTRL+9 viruses in biological replicates for RNA-seq analysis at days three (n=2 per condition) and six post infection (n=3 per condition; **Figure S4A-S4D**).

RNA-seq showed that viral RNA (vRNA) was transcribed at similar levels between PB1-UFO mutant and control viruses (**Figure S4A**), consistent with the minimal difference in viral lung titers (**Figure 4C, top panels**). Importantly, mutant and control viruses exhibited a distinct transcriptome signature at day six but not day three post infection (**Figure S4B and S4C**), as observed through differential gene expression analysis (**Figure S4C**). Our analysis also suggested that the two PB1-UFO mutant viruses behaved similarly to each other during infection (**Figure S4B**; Comparison m3 v p9), supporting the conclusion that the difference in gene expression between control and mutant viruses are due to loss of PB1-UFO, and not just alterations in viral RNA sequences. Similar trends in gene expression differences are present in the top 32 differentially expressed genes (**Figure S4C**). Genes differentially expressed at day 6 post infection are predominantly related to angiogenesis and protein folding (**Figure S4D**).

To check whether PB1-UFO expression could be detected by the immune system, we inserted the SIINFEKL (SIIN) model MHC class I peptide (SIIN) into the 5’ UTR upstream of the native PB1 initiation codon and in frame with PB1-UFO in a recombinant PR8 virus (**Figure 4D**). SIIN is efficiently processed by the class I pathway and generates a high affinity complex with the mouse Kb class I molecule that can be detected on the cell surface with high specificity and sensitivity by the 25-D1.16 mAb (Porgador et al., 1997) (**Figure S3C**). HEK293Kb cells infected with the PB1-UFO (SIIN) virus demonstrated increased staining in flow cytometry relative to control uninfected cells or cells infected with wildtype PR8 virus (**Figure S3D**). Extending this finding, mouse DC2.4 cells infected with PB1-UFO (SIIN) PR8 activated transgenic OT-I CD8+ T cells (highly specific for Kb-SIIN (Hogquist et al., 1994)) as determined by upregulation of CD25 and CD69 and also induced OT-I T-cell proliferation when compared to the negative controls. (**Figure 4E**). As a positive control, we used a recombinant IAV expressing SIIN(PB1-Ub-SIIN) at high levels (Wei et al., 2019). These data confirm that PB1-UFO is translated, expressed during infection and that T cell immunosurveillance extends to peptides encoded by uvORFs.

### Conservation of host-IAV proteins across strains and time

To establish the contribution of the PB1-UFO to viral fitness we examined its conservation among all H1N1, H3N2 and H5N1 IAV strains deposited in public sequence databases (Shu and McCauley, 2017). Due to its high mutation rate, IAV evolution occurs extremely rapidly, and conservation of the ORF provides strong evidence for its contribution to IAV transmission in its natural hosts. The PB1-UFO is highly conserved across these three virus subtypes, all of which encode proteins of similar length and amino acid composition (**Figure 5A**). Truncating mutations occurred relatively infrequently, at 8% of H1N1 sequences, 3% of H5N1 sequences, and not present in H3N2 isolates.

**Figure 5.**
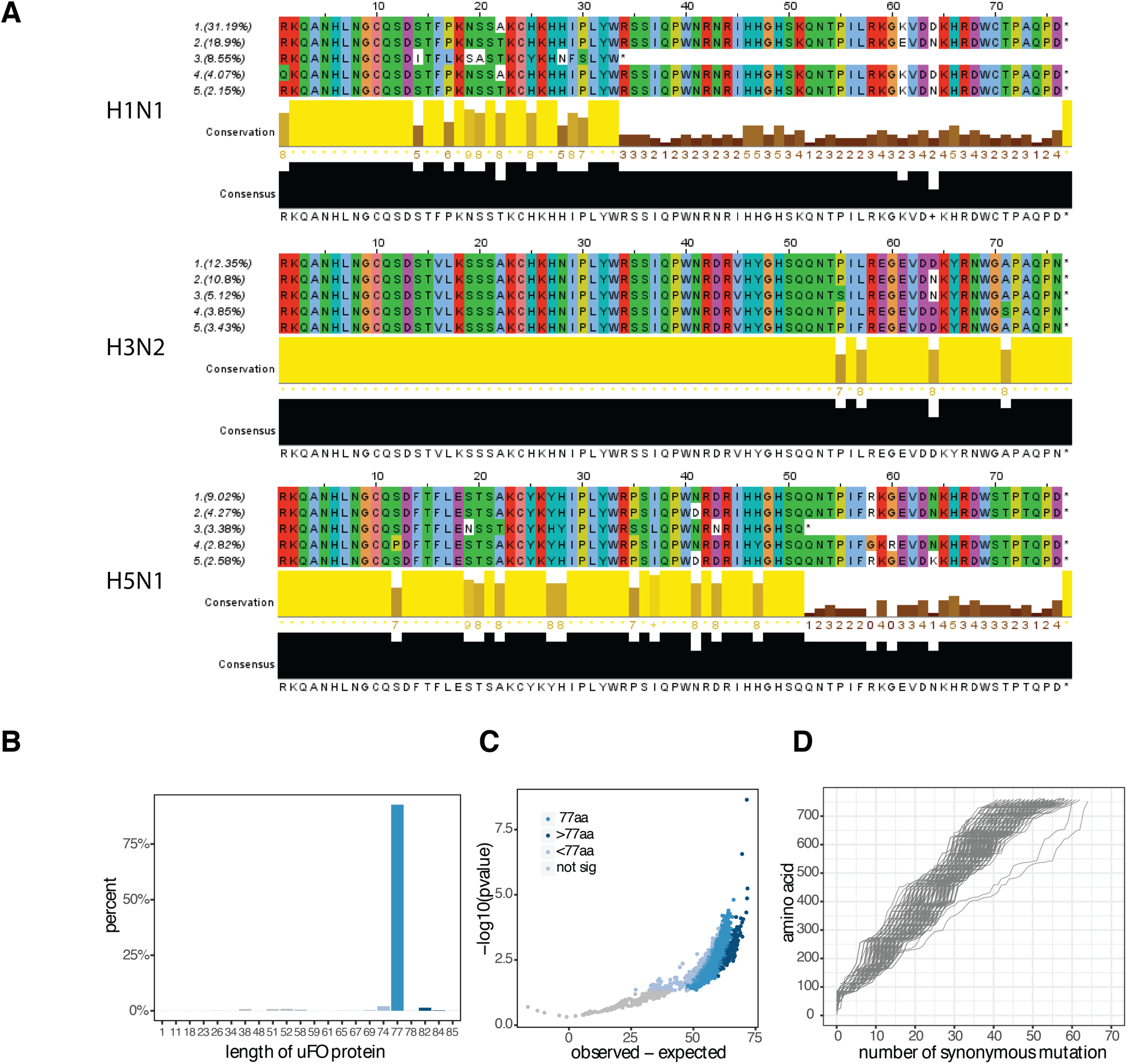
Bioinformatics analysis on conservation of PB1-UFO protein sequences. (A) Top five most common PB1-UFO protein sequences in three Influenza A strains, H1N1, H3N2 and H5N1. (B) Density plot of predicted length of H3N2 PB1-UFO protein sequences. Over 92% of sequences are predicted to generate a protein of 77aa (medium blue), ∼3% are shorter than 77aa (light blue), ∼1.5% are longer than 77aa (dark blue), rest of sequences are predicted not to generate PB1-UFO protein (grey). (C) P value distribution/volcano plot of H3N2 PB1-UFO protein sequence length. Each dot represents the difference between observed length and expected length of each individual sequence. (D) The line plot shows the number of synonymous mutations in frame of WT H3N2 PB1 (x-axis) that mutate stop codons in frame of H3N2 PB1-UFO (y-axis).

We next designed a statistical model (**Figure S5A**) to query whether the conserved length of PB1-UFO occurs more frequently than expected by chance. To increase statistical power, we focused only on sequences derived from the H3N2 subtype of viruses as they had the most abundant number of full-length, unique PB1 segment sequences. Over 92% of H3N2 PB1-segment sequences encode a 77-amino acid PB1-UFO (**Figure 5B and 5C**). This is highly significant given that the random mutation model predicts an average ORF of ∼19 amino acids (**Figure S5B**). Likewise, analyses on synonymous mutations showed that PB1-UFO is highly likely to maintain a long amino acid sequence pattern, implying that the maintenance of longer sequences is due to protein function rather than the random production of short peptides (**Figure 5D**).

### Evolutionary analyses on chimeric protein maintenance

In an effort to quantify and infer selection of PB1-UFO in H3N2 strain of influenza virus over time we used a frequency propagator model (Luksza and Lassig, 2014; Strelkowa and Lässig, 2012) (**Figure 6A–6B**). We compared the likelihoods of non-synonymous and/or synonymous mutations to reach fixation in the PB1-UFO coding sequence of the IAV 5’ UTR (R1, **Figure 6C**) to the corresponding likelihood of synonymous mutations, which should evolve at near neutrality, occurring in the main PB1 coding sequence (R3, **Figure 6C**). A similar analysis was done using the nucleotide sequences where PB1-UFO and PB1 ORF overlap (R2, **Figure 6D**). Our analysis suggests that nucleotide mutations are overall strongly repressed in the IAV 5’UTR, consistent with its role in priming viral transcription and viral packaging (**Figure 6C**, Black line). Despite this, the fixation probabilities of synonymous mutations occurring in PB1-UFO (**Figure 6C**, red line; Frequency propagator ratio =0.263+/-0.094) were 2 fold increased over that of non-synonymous mutations (**Figure 6C**, blue line; Frequency propagator ratio =0.134+/-0.040). This suggests that mutations that preserved the PB1-UFO peptide sequence are better tolerated within the viral UTR. In contrast, when the nucleotide sequence of PB1-UFO overlapped PB1 (R2, **Figure 6D**), there was no difference in fixation rates between synonymous or non-synonymous mutations (Frequency propagator ratio = 0.924 +/-0.650 (synonymous; red lines) and 1.121+/-0.225 (non-synonymous; blue lines) (**Figure 6D**), indicating that changes to amino acid sequence are more tolerated in the C-terminal of PB1-UFO. Taken together, our analyses suggest that while selection is heterogeneous across the PB1-UFO frame, there is a positive selection pressure to maintain both the PB1-UFO protein length, and N-terminal sequences.

**Figure 6.**
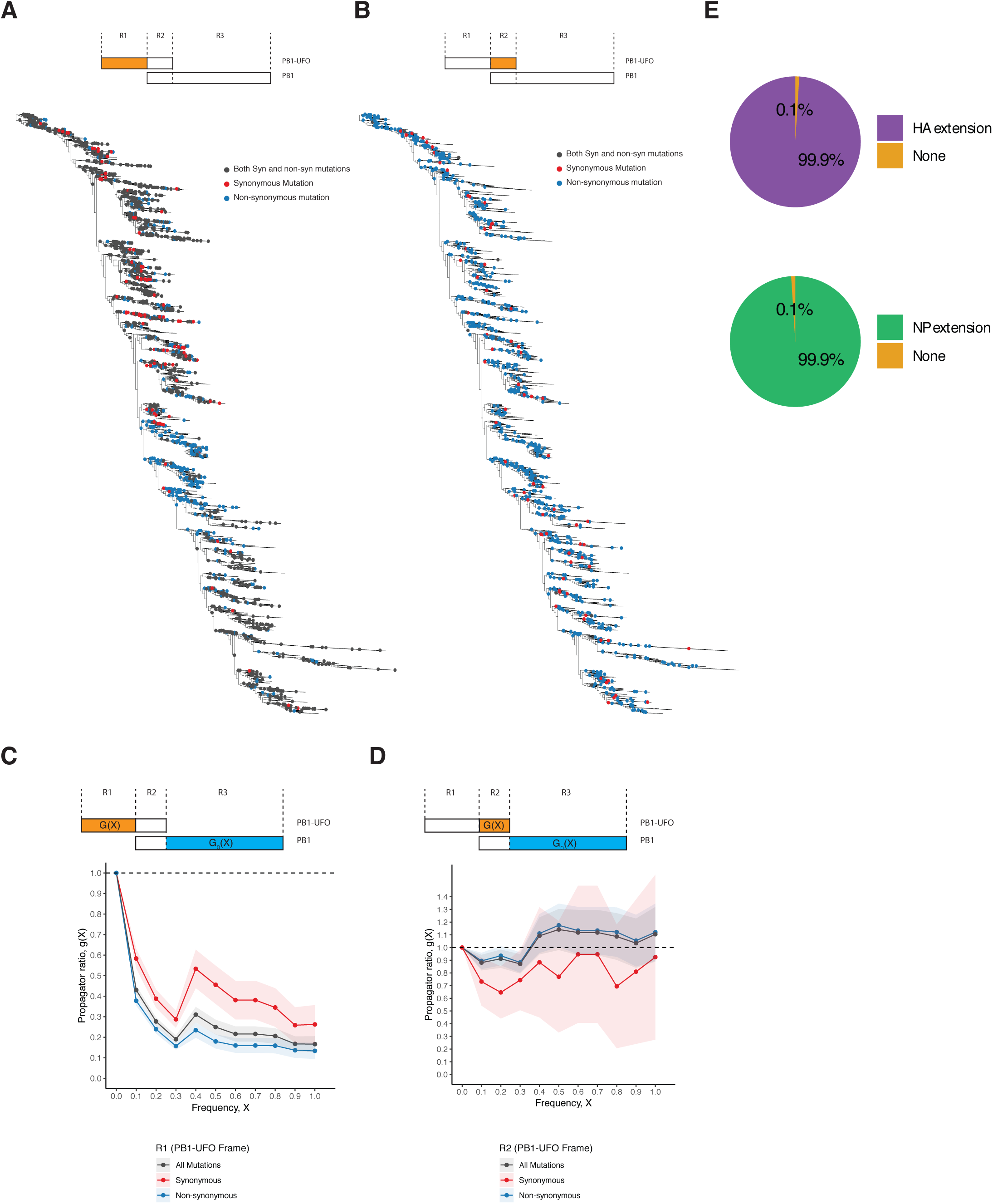
Evolutionary analysis on H3N2 PB1-UFO protein sequences. (A) Strain tree of H3N2 IAV viruses. Mutations occurring in the N-terminal PB1-UFO frame overlapping the viral 5’UTR (region1, R1, top panel; yellow region) are indicated as color dots. (B) Same as in A, but for mutations occurring in the C-terminal PB1-UFO frame overlapping the PB1 (region 2, R2, top panel; yellow region). (C) Frequency propagator ratio of the indicated classes of mutations occurring in the N-terminal PB1-UFO frame overlapping the viral 5’UTR (region1, R1; yellow). Fixation probabilities were compared to those of synonymous mutations occurring in the region of PB1 that does not overlap PB1-UFO (region 3, R3; blue). Error bars indicate sampling uncertainties. g(X) <1: negative selection, g(X) ≈1:weak/heterogeneous selection; g(X) > 1: positive selection. (D) Same as in (C), but for C-terminal sequences of PB1-UFO frame overlapping the PB1-frame (region 2, R2; yellow). (E) Percentage of observed HA and NP N-terminal extension protein sequences.

Similarly, HA- and NA-UFO extensions are conserved more than 99% of H3N2 and are under positive selection (**Figure 6E**). Overall, our result indicates that mutations that disrupt the amino acid sequences of PB1-UFO and/or viral extensions are not well tolerated. IAV thus maintains the capacity, throughout strains and time, to encode for chimeric proteins. This indicates a role for such host-dependent viral protein diversity in viral tropism and life cycle.

## DISCUSSION

In this manuscript, we describe the existence of a novel mechanism employed by IAV to generate hitherto uncharacterized host-virus chimeric proteins. This mechanism employs the generation of host-virus chimeric RNAs that are translated into chimeric proteins. We show that during IAV infection, two classes of chimeric proteins are made: (1) viral proteins with host-encoded N-terminal, and (2) chimeric host-virus proteins with novel open reading frames, which we termed uvORF proteins. We show that these gene products are expressed in infected cells, surveilled by CD8+ T cells and modulate the antiviral response.

### Chimeric UFO proteins: Novel proteins and N-terminal extensions

In human genes, there is increasing evidence that upstream start codons (uAUGs) in the 5’ UTR initiate translation of short ORFs (Calvo et al., 2009; Wang and Rothnagel, 2004). uAUGs/uORFs are thought to be mainly important in regulating expression of downstream ORFs by controlling ribosomal scanning efficiencies (Calvo et al., 2009). However, there is evidence that suggests that some uORFs encode biologically active peptides that contribute to evolutionary fitness (Andrews and Rothnagel, 2014; Combier et al., 2008; Wen et al., 2009).

We have characterized the N-terminal HA and NP extensions, as well as PB1-UFO, because they could be identified unambiguously by analyzing chimeric host-virus or UTR derived peptide through mass spectrometry (whose sequence do not exist in ‘conventional’ human and viral proteome databases). The expression of other uvORF proteins remains to be determined. It is important to recognize that other uvORFs or N terminal extensions, like the chimeric HA and NP described here, might be difficult to detect due to their N-term heterogeneity and partial overlapping sequences with the canonical protein.

In fact, we showed that based on the length of host snatched sequences and the viral UTRs, the chimeric-protein extension bear N-termini consisting essentially of hypervariable peptides encoded by host-derived RNA. In the cell, proteins containing variable sequences (“quasi protein-species”) can be generated in expressed proteins under normal conditions. This occurs through natural errors in protein synthesis as the translation apparatus is tuned to optimize the occurrence of semi-random amino acid substitutions. Translational fidelity is adaptive, maintained by cell and tissue type, and likely functions to cushion stress (Ribas de Pouplana et al., 2014). For instance, conditions of oxidative stress alters the specificity of Methionine(Met)-amino acyl synthetase, increasing Met-charging of non-Met tRNA to increase the Met content of proteins (Netzer et al., 2009). This presumably protects them from oxidative damage (Levine et al., 1996). Instead of relying wholly on adaptive mis-translation, IAV uvORFs appear to have a built-in mechanism to diversify their proteome during infection. One intriguing hypothesis is that usage of human-derived protein appendices might ‘confound’ MHC-class I surveillance.

In a similar vein, uORFs are particularly common in host cell mRNAs encoding regulatory and stress-responsive proteins (Bondke Persson et al., 2015; Starck et al., 2016; Young and Wek, 2016), suggesting that these genetic elements respond to changes in the cell’s environment. Stress, leads to global translational repression and preferential usage of uAUGs. This results in pervasive translation of human uORFs as documented in cancer cells (Sendoel et al., 2017) and activated T cells (Starck et al., 2016). In line with this, our data indicate that uAUGs are particularly abundant in high expressing genes that are cap-snatched by IAV. By generating human-viral mRNA chimeras during infection IAV may be co-opting the altered host mRNA expression to drive the expression of its newly expanded proteome.

### Evolutionary considerations: Overprinting and the mis-naming of UTRs

Genetic overprinting typically occurs when a pre-existing reading frame acquires mutations that enable translation in alternative reading frames while maintaining function of the ancestral frame. This is an important mechanism to create new proteins, especially in the context of compact genomes (viral, prokaryotic, eukaryotic organelles) with little coding capacity (Keese and Gibbs, 1992; Kovacs et al., 2010; Poulin et al., 2003; Sabath et al., 2012).

Alternative reading frames created by overprinting can be translated by two mechanisms. One way is via leaky scanning ribosomes that bypass the canonical AUG and decode a downstream out-of-frame initiation codon. Important viral virulence factors, like PB1-F2 from influenza virus, are generated by such a mechanism (Chen et al., 2001). Alternative reading frames may also be translated via ribosome frameshift, in which ribosomes slip and skips (either forward or backward) one or two nucleotides to shift to a new reading frame. IAV uses this process to create PA-X (Jagger et al., 2012). HIV also uses this process to express and regulate the expression of Gag and Gag-Pol proteins, which are encoded by the same ORFs (Fernandes et al., 2016).

While genetic overprinting could be selectively advantageous for some organisms, maintaining overlapping ORFs requires additional regulatory mechanisms (e.g. regulating dynamic expression of multiple proteins upon stimulation and/or ways to stop expression of one ORF in favor of the second). These limitations, along with the fact that a mutation in one ORF will also often affect a second ORF, ends up imposing too many constrains, thus limiting the functional evolutionary space that pathogens require to sample as a mean to adapt to hosts.

PB1-UFO represents a unique product of overprinting because it is encoded by sequences from two organisms: virus and host, with host sequences providing translatability to viral UTR sequences. Our analyses suggest that PB1-UFO is undergoing stabilizing selection in the 5’UTR, where divergent forms of the protein, generated by non-synonymous mutations, appear to be preferentially removed from the population. This implies that PB1-UFO support viral fitness, as we would not otherwise expect differences in fixation probabilities of synonymous or non-synonymous mutations occurring in the IAV 5’UTR.

### A new player in the host-pathogen arms race

The capacity of a pathogen to overcome host barriers and establish infection is based on the expression of pathogen-derived proteins. To understand how a pathogen antagonizes the host and establishes infection we need to have a clear understanding of what protein a pathogen encodes, how they function, and the manner in which they contribute to virulence. The current dogma about many life-threatening pathogens is that they encode a small, finite number of proteins because of their limited genomes. RNA viruses, including IAV, are a prime example of this paradigm. We now show that there is another level of complexity to this equation.

## AUTHOR CONTRIBUTIONS

Conceptualization, A.G.-S., J.W.Y. and I.M.; Methodology, I.M., J.W.Y., Y.M., M.A., G.W., J.H.; Formal Analysis, Y.M., M.A., G.W., J.H., N.Z., J.N., N.M., J.G., J.W., J.J., M.C., Z.P., H.v.B., M.L., E.R.M.; Investigation, Y.M., M.A., G.W., J.H., N.Z., J.N., N.M., J.J., M.C., Z.P., H.v.B., M.L., E.R.M.; Resources, Y.M., M.A., J.J., M.C., H.v.B., E.R.M., A.G.-S.; Writing – Original Draft, I.M., J.W.Y., Writing – Review & Editing, I.M., J.W.Y., Y.M., J.H., M.A., G.W., H.v.B., M.L., B.D.G, E.R.M, A.G.-S.; Visualization, Y.M., M.A., G.W., J.H., J.J., M.C., Z.P., E.R.M; Funding Acquisition, A.G.-S, I.M.; Supervision, I.M.

## ACKNOWLEDGMENTS

We thank the Genomics and Mouse facility at Icahn School of Medicine at Mount Sinai, the Global Health and Emerging Pathogens Institute (GHEPI) at Mount Sinai, and the entire Marazzi Lab team. I.M. is supported by Burroughs Wellcome Fund 1017892 and by Chan Zuckerberg Initiative 2018-191895. I.M. and H.v.B. are supported by NIH grant R01AI113186. A.G.-S. and I.M. are supported by the NIH grant U19AI135972 FLUOMICS.

## MATERIALS AND METHODS

### Cells

Human embryonic kidney 293T cells, Madin-Darby canine kidney (MDCK) cells, and Human lung carcinoma epithelial A549 cells were maintained in Dulbecco’s modified Eagle’s medium (DMEM; Corning) containing 10% newborn calf serum (FBS; Peak Serum) and antimicrobial drugs. Human. All cells were maintained at 37 °C with 5% CO2.

### Viruses

Using plasmid-based reverse genetics (Fodor et al., 1999), we generated recombinant influenza viruses with PB1 mutations by using the A/Puerto Rico/8/1934 (PR8) strain as the backbone. A wild-type (WT) recombinant (PR8 WT) was generated, as well as four PB1 substitution mutants.

The first mutant, bearing a premature stop codon in the PB1-UFO protein at the position of three nucleotides before the start of PB1 open reading frame, was named as (PB1-UFO KO-3). The second mutant, which preserved expression of full length PB1-UFO protein even with a point mutation at the position of three nucleotides before the start of PB1 open reading frame, was named as (PB1-UFO Ctrl-3). This virus acted as a control of PB1-UFO KO-3. The third mutant containing a stop codon in the PB1-UFO protein at the position of nine nucleotides after the start of PB1 open reading frame was named as (PB1-UFO KO+9). The fourth mutant, which preserved the expression of full length PB1-UFO protein even with a point mutation at the position of nine nucleotides after the start of PB1 open reading frame was named as (PB1-UFO Ctrl+9). This virus acted as a control of PB1-UFO KO+9. Mutations were confirmed by sequencing both plasmids and viruses. The stock virus titers were the average of three independent experiments.

### Growth kinetics of Viruses in Cell Culture

MDCK cells were infected with viruses at a multiplicity of infection (MOI) of 0.001, incubated for one hour at 37 °C, washed twice, and then cultured with Opti-MEM and TPCK-treated trypsin at 40°C and 37°C for 72 h. Supernatants were collected at the indicated time points. Hemagglutination titer (HA) were tested in 0.5% turkey red blood cells and virus titers were determined by plaque assay in MDCK cells.

### Mouse studies

All mice procedures were performed following protocols approved by the Icahn School of Medicine at Mount Sinai Institutional Animal Care and Use Committee (IACUC). All the animal studies were carried out in strict accordance with the recommendations in the Guide for the Care and Use of Laboratory Animals of the National Research Council. Eight-week-old female BALB/c mice were obtained from Jackson Laboratories (Bar Harbor, ME). Mice were anesthetized by intraperitoneal injection of a mixture of ketamine and xylazine before infection.

Groups of five mice were inoculated intranasally. with 100, 50, 25, 10, or 5 PFU of virus. Mice were monitored daily for clinical signs of illness and weight loss. Upon reaching 75% of initial body weight, animals were humanely euthanized with carbon dioxide (CO2) as per the IACUC protocol.

Groups of five mice were intranasally (i.n.) infected with 100 plaque-forming unites (PFU) of viruses in a volume of 50 µl, two and three mice were euthanized on 3 and 6 days post-inoculation (d.p.i.), respectively. The middle lobe of the lung was collected for total RNA extraction, and the post-caval lobes of the lung was collected to determine virus titers by plaque assay on MDCK cells.

### RNA sequencing

After adaptor removal with cutadapt (Martin, 2011) and base-quality trimming to remove 3′ read sequences if more than 20 bases with Q <20 were present, paired-end reads were mapped to the mouse (mm10) reference genome with STAR (Dobin et al., 2013), and gene-count summaries were generated with featureCounts (Liao et al., 2013). DESeq2 (Love et al., 2014) was used to variance-normalize the data before a 2-factor model (gene ∼ ConditionTime + Mutant) was applied to identify differentially expressed genes. RNA-seq raw data are deposited in GEO under accession GSE128519.

### Proteomic Strategy

Mass spectrometry was performed using purified lysates obtained from PR8 IAV infected A549 cells. Targeted identification of chimeric proteins was conducted using datasets derived from the entire human and IAV reference sequence merged with the set of predicted IAV uvORFs and viral protein extensions. Common contaminants were filtered out and missing values in the data matrix were attributed an intensity score of 0.

### Ribo-seq analysis

Ribosome footprint reads were trimmed with cutadapt (Martin, 2011), and aligned to the human (hg38) and A/Puerto Rico/8/1938 (H1N1) genomes with STAR (Dobin et al., 2013). The 5’ end mapping was then performed for all reads aligning to the influenza genome. Host-derived transcriptional primer sequences were extracted from reads with partially mapping to the 5’ end of each segment. Analysis of AUG composition was performed using custom in-house Perl scripts which are available upon request.

### Antigen Expression and T cell immunosurveillance Assays

HEK293T cells stably expressing mouse K^b^ MHC-I (HEK293K^b^) were infected with influenza A viruses. At 18 hours post infection, cells were stained with Alexa 647-labelled MAb 25D-1.16 (anti-K^b^-SIIN) to measure surface expression of K^b^-SIIN complexes flow cytometry. For T-cell activation assays, OT-I T-cells were harvested from the spleen and lymph nodes of OT-1 transgenic mice and purified on the AutoMACS with the CD8a+ T Cell Isolation Kit (Milteny, Germany), and stained with CellTrace Violet (Thermo Fisher, Waltham, MA) DC2.4 cells were infected with influenza A viruses for 18 hours, and then co-cultured with OT-I T-cells. T-cells were stained with anti-CD25 and anti-CD28 labeled antibodies at 24 hours post co-culture for activation assays. T-cell proliferation assays were conducted at 48 hours post infection by measuring CellTrace Violet staining by flow cytometry.

#### Computational analyses

##### Sequence data set

Our study is based on a data set of 26,472 human influenza A/H3N2 sequences available from the GISAID database (Shu and McCauley, 2017), which contain 6,244 unique PB1 strains. We included only full length sequences using a custom script that is available upon request.

##### Random sequence model

We constructed codon usage matrix for each of individual nucleotide sequence. Using the codon usage table, a protein sequence in open reading frame is used as input to generate multiple random nucleotide sequences. We then translate random nucleotide sequences to protein sequences in frame which may generate PB1-UFO protein. Using a custom script, we calculate the average stop codon positions of random PB1-UFO protein sequences as its expected value. By comparing with its expected value, we determine the likelihood that the translated PB1-UFO sequence was obtained randomly and its deviation from the expected PB1-UFO length.

##### Strain tree reconstruction

Our analysis is based on an ensemble of strain trees obtained from the PB1 sequence data set. Such trees describe the genealogy of influenza strains resulting from an evolutionary process under selection(Strelkowa and Lässig, 2012). The tree ensemble is obtained with FastTree (Price et al., 2010), which very time-efficiently reconstructs maximum-likelihood phylogenies. We use a general time-reversible model. We refine the tree topology with RAxML(Stamatakis, 2014). Given the output topology, we reconstruct maximum-likelihood sequences and timing of internal nodes with the TreeTime package (Sagulenko et al., 2018).

##### Mapping of mutations

Maximum likelihood maps point mutations between directly related strains onto the branches of the tree. A mutation on a given branch marks an origination event of a single-nucleotide polymorphism, that is, the appearance of a nucleotide difference between the strains descending from that branch and its ancestral lineage. A reconstructed strain tree with all mapped mutations, which are partitioned into two classes: (a) synonymous mutations, (b) nonsynonymous mutations. These mutations are the basis of our fitness model (Luksza and Lassig, 2014).

##### Frequency propagator ratio analysis

Our analysis is based on a set of codons in PB1-UFO coding sequence overlapping the IAV 5’UTR (R1), the overlapping sequence between PB1-UFO and PB1 ORF (R2), and those in the main PB1 coding sequence (R3) respectively. To quantify selection on a class of mutations, we use the frequency propagator ratio

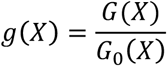

where G(X) is the likelihood that a mutation in a given class reaches frequency X, and G_0_(X) is the likelihood for synonymous mutations occurring in the main PB1 coding sequence, which should evolve near neutrality. To predict the evolutionary direction of a given subset of codons, we compare fixation probabilities (or probabilities of reaching high frequencies) of mutations in that region with those in the null-class region R3.

## FIGURE LEGENDS

**Figure S1.**
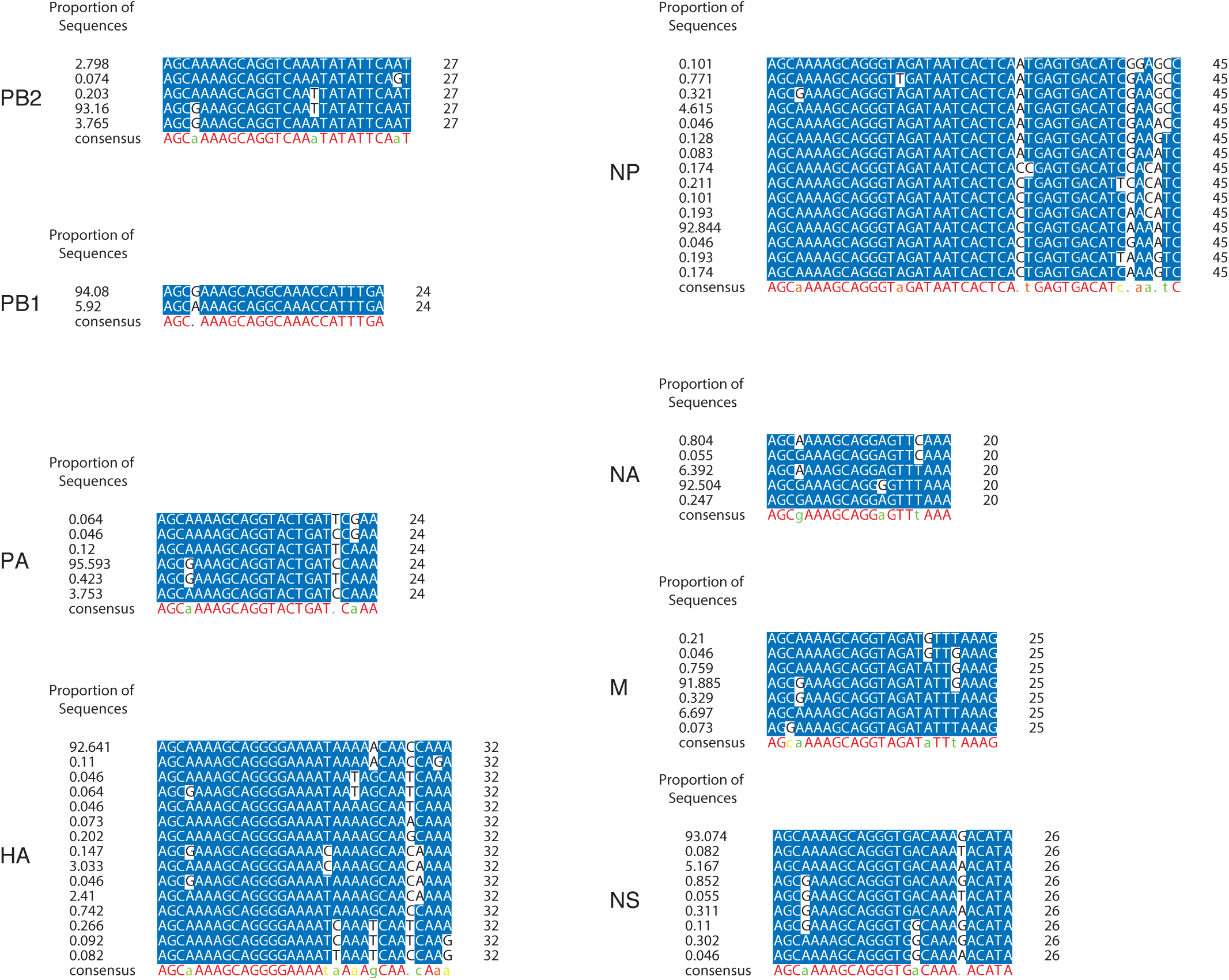
Viral 5’UTRs are conserved (Related to Figure 1). Multiple sequence alignments of H1N1 IAV 5’UTRs per segment. The overall distribution of each unique nucleotide sequence is indicated on the left, and the consensus sequence of each UTR is indicated below each alignment.

**Figure S2.**
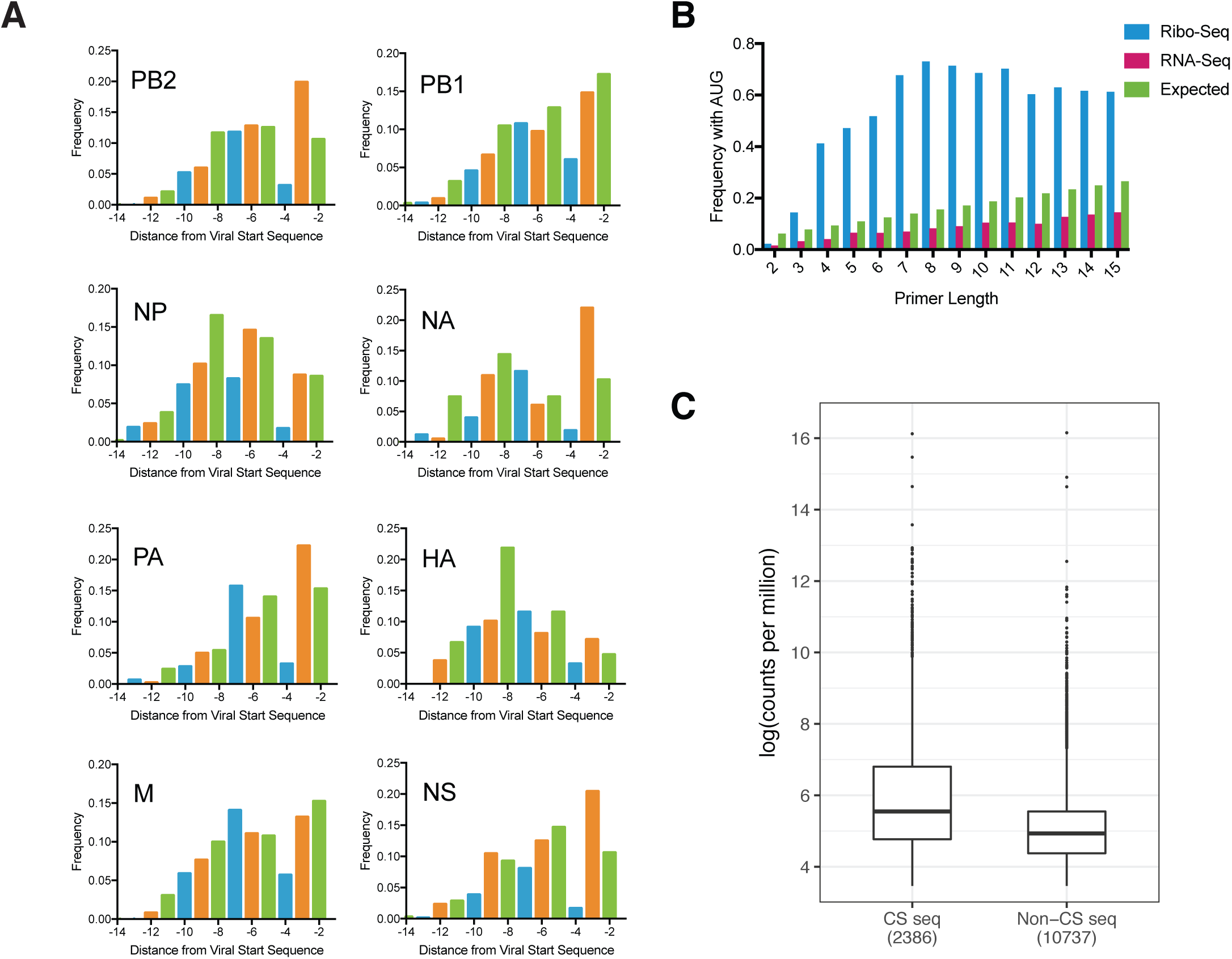
IAV 5’UTRs are associated with active ribosomes (Related to Figure 2). (A) Frequency of AUG codons by position relative to the viral transcription initiation site. (B) Frequency of AUGs at each position in primers compared to length for harringtonine-arrested ribo-seq and RNA-seq datasets. Expected frequency shown in green. (C) Log (counts per million) of transcripts that are cap-snatched by IAV or control transcripts.

**Figure S3.**
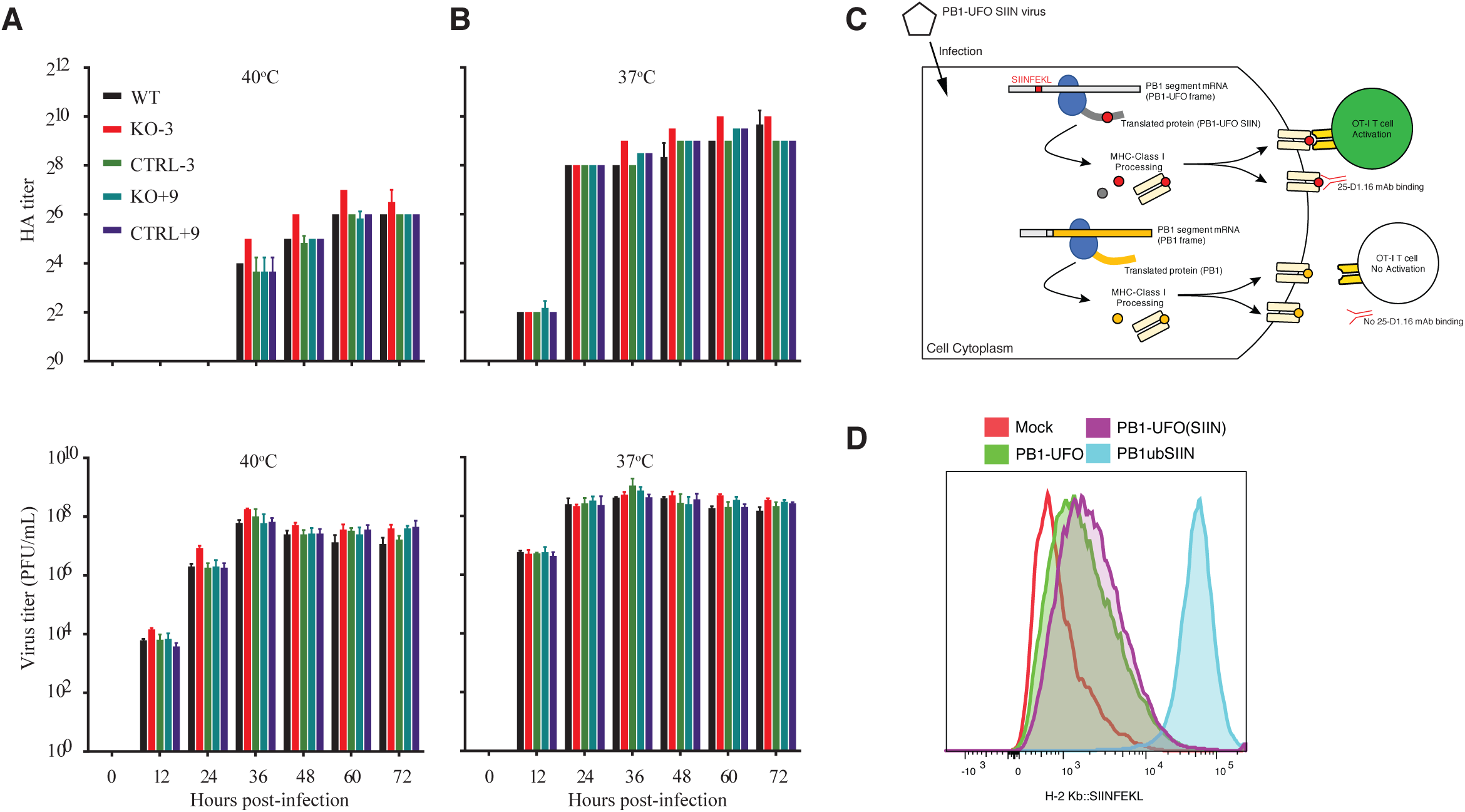
Controls for Figure 4 (Related to Figure 4). (A) and (B) shows growth properties of viruses in MDCK cells. Cells were infected with viruses at MOI of 0.001 and incubated at 40°C (A) and 37°C (B). (C) Schematic of SIINFEKL mechanism of action. (D) SIINFEKL expression from 293Kb cells infected with PB1-UFO (SIIN) virus.

**Figure S4.**
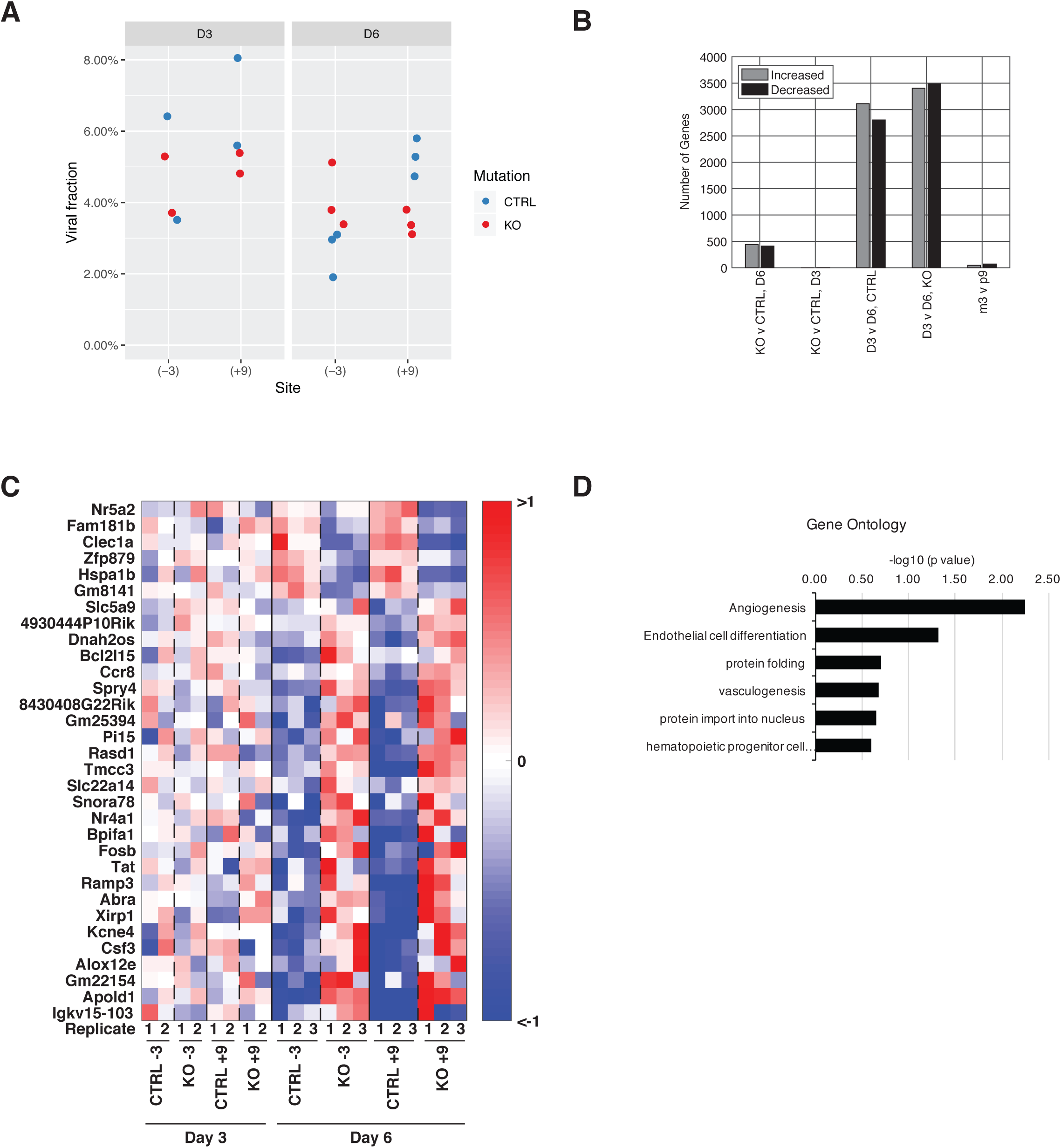
RNA-Seq analysis on PB1-UFO mutant and control virus infected mice (Related to Figure 4). (A) Viral RNA levels in PB1-UFO mutant and control viruses in the lungs of infected mice at days 3 and days 6 post infection. (B) Two factor model analyses of RNA sequencing data of PB1-UFO mutant and control viruses infected mouse lungs at days 3 and days 6 post infection. (C) Heatmap showing the top 32 differentially expressed genes (FDR < 0.1, |Log2FC| > 1) when comparing PB1-UFO mutant and control virus infected lungs at day 6 post infection. (D) Gene ontology of genes predicted to be differentially expressed during infection of control or PB1-UFO deficient viruses in mice.

**Figure S5.**
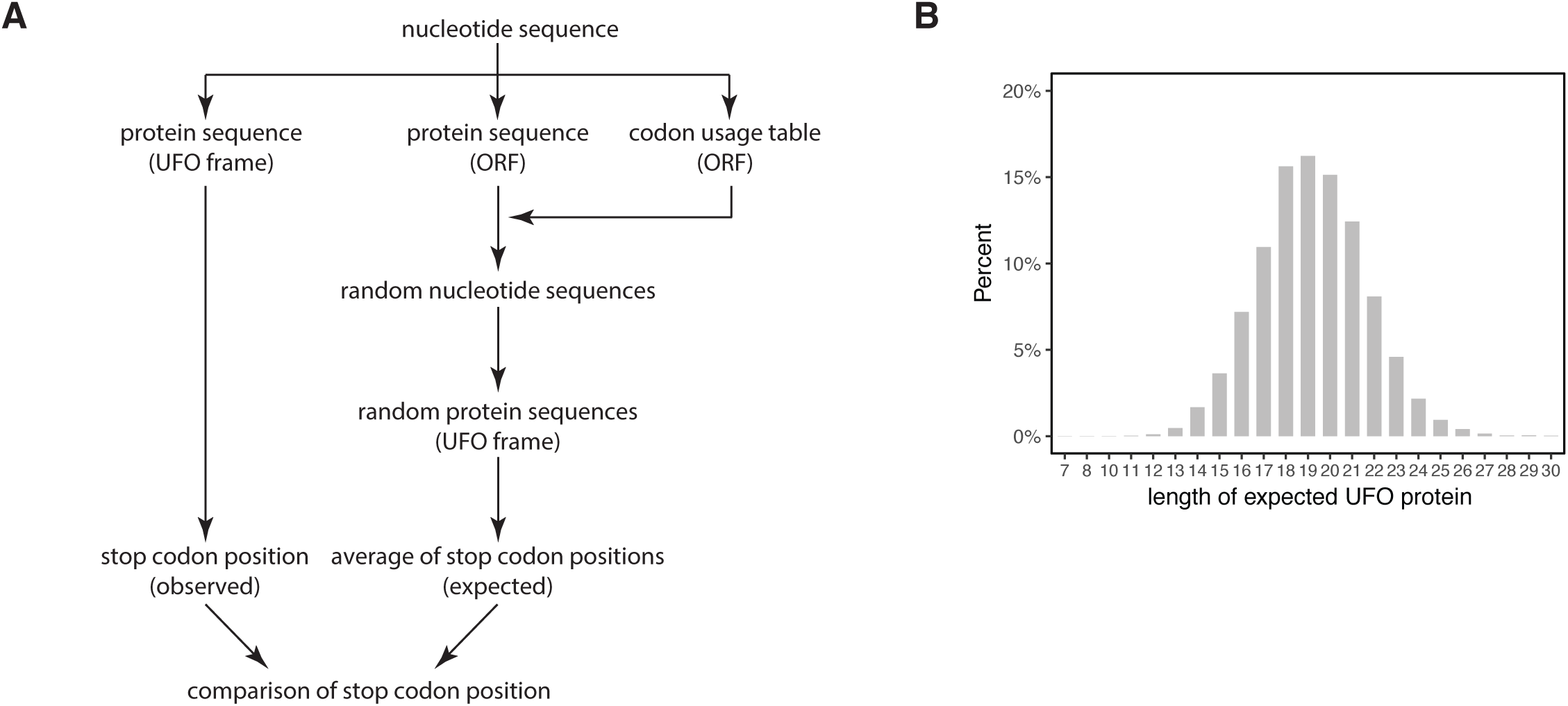
Controls related to Figure 5 (Related to Figure 5). (A) Schematic of predicted sequence length model of PB1-UFO proteins. (B) Density plot showing the expected lengths of H3N2 PB1-UFO proteins, based on random codon-shuffled sequences.

